# Overcoming Cisplatin Resistance in TP53-null Colon Cancer Organoids

**DOI:** 10.1101/2025.02.12.637569

**Authors:** Sana Khalili, Paige Heine, Minou Khazan, Carolyn E. Banister, Sidney E. Morrison, Phillip J. Buckhaults

**Affiliations:** University of South Carolina, Columbia SC; Medical University of South Carolina, Columbia, SC

## Abstract

**Cisplatin chemotherapy of colorectal cancer (CRC) is associated with dose-limiting side effects and the development of drug resistance, resulting in reduced therapeutic effectiveness. The resistant phenotype in colon cancer is primarily due to changes in p53-regulated DNA damage signaling and /or defects in the cellular mismatch-repair pathway. Therefore, enhancing the efficacy of cisplatin chemotherapy remains a significant challenge. In this study, we used a TP53-KO patient-derived colon tumor organoid model to perform a genome-wide CRISPR KO screen in the absence and presence of cisplatin and identified gene knockouts that re-sensitize cisplatin-resistant TP53-KO colon cancer organoids to cisplatin treatment. Knockout of genes in the DNA Repair pathways, including Fanconi Anemia (FA cause re-sensitization of TP53-KO colon cancer cells to cisplatin. Inhibition of genes ERCC6, FANCL, and BRIP1 enhances cisplatin-induced cell death in TP53-KO colon cancer organoids. These findings suggest that targeting these pathways could be an effective approach to overcome chemoresistance of TP53-muatnt colon cancer cells to cisplatin.**

## Main

Colorectal Cancer (CRC) ranks as a leading cause of cancer-related deaths worldwide ^1^. Cisplatin (CDDP), a platinum-based compound, is widely used as an anti-cancer agent across various malignancies, effectively targeting cancer cells by inducing DNA damage and causing cell cycle arrest and apoptosis ^2^. Despite its initial efficacy, the effectiveness of cisplatin is progressively compromised in colon cancer cases duo to the development of resistance, resulting in tumor relapse and therapeutic failures ^3^. Extensive studies has identified that these resistance mechanisms are linked to enhanced DNA repair and activation of alternative apoptotic pathways ^3^. Therefore, targeting these pathways could significantly mitigate the chemotherapeutic resistance observed with Cisplatin.

The majority of colon cancers are initiated by the loss of function of tumor suppressor genes such as APC, MMR genes and TP53. The transcription factor p53 is pivotal in mediating the apoptotic response of tumor cells to therapeutic agents ^4^. Mutations in the TP53 gene are primarily responsible for disruptions in apoptotic processes ^5 6^. Both *in vitro* and clinical studies have showed that stabilizing and activating wild-type p53 is essential for apoptosis triggered by cisplatin ^7 8^. Therefore, mutations in TP53 can contribute to cisplatin resistance by impairing the apoptotic response necessary for its therapeutic efficacy ^9^.

Genome-wide CRISPR knockout (KO) screening in human cell lines can reveal dependencies that could be therapeutically exploited ^10^. CRISPR/Cas9 knockout screening on two-dimensional model systems provides valuable insights into genetic dependency for tumor cell lines ^11 12^. CRISPR/Cas9 knockout screening on three-dimensional models ^13^ may reveal additional insights not apparent in 2D cell lines. Organoids are self-organizing mammalian adult stem cells and are strong tools for *ex vivo* tissue morphogenesis and organogenesis simulations ^14^. Using these organoids in CRISPR/Cas9 knockout screens can reveal three-way synthetic lethal interactions between TP53 mutations, DNA-damaging agents, and secondary gene mutations. Our previous study on human embryonic stem cells (hESC) shows that genome-wide CRISPR KO library screening can unveil targets for resensitizing chemotherapy-resistant stem cells in presence and absence of DNA damaging agents ^15^. These interactions pinpoint secondary genes that can be identified through CRISPR KO library screenings of human colon cancer organoids, in presence and absence of DNA damaging agents such as Cisplatin. This approach could identify novel therapeutic targets for resensitizing chemotherapy-resistant cells, leading to therapeutic strategies tailored to overcome drug resistance in colon cancer.

In our previous study, we created isogenic pairs of TP53-wildtype (TP53-WT) and TP53-knockout (TP53-KO) human colon tumor organoids (F147T) and demonstrated that TP53-deficiency induces resistance to cisplatin in these colon cancer organoids ^16^. Building on this model, we conducted a genome-wide CRISPR dropout library screen (Brunello) ^17 18^ using these isogenic pairs of colon cancer organoids to identify colon cancer dependencies ^16^. We cultured those organoids, screened with genome-wide CRISPR KO library, for 58 days and observed dropout of essential genes, including DepMap common essentials (19%) (https://depmap.org), due to their necessity for cell survival. We then continued culturing these cisplatin-resistant TP53-KO organoids in the presence and absence of cisplatin for an additional 28 days (total of 86 days in the culture). We identified and validated several novel cisplatin resensitizing targets involved in DNA repair pathways. These findings support the idea of inhibition of Fanconi anemia and other DNA repair genes as a potential chemo-sensitization to cisplatin in resistant-TP53 mutant colon cancer in combinational therapy.

## Results

### TP53 loss causes resistance to Cisplatin

Following DNA damage by cisplatin, cells initiate a DNA-damage response that can lead to either the repair of the lesion, thereby promoting resistance to treatment, or cell death via the activation of apoptotic pathways. Many clinical studies have underscored the prognostic significance of the human tumor suppressor protein p53 across various types of human cancers. Often, the overexpression of mutated p53, which has reduced or lost functionally, is linked to resistance to standard treatments ^6 7 19 20^. This includes resistance to drugs such as cisplatin, alkylating agents (temozolomide), anthracyclines (doxorubicin), antimetabolites (gemcitabine), anti-estrogen (tamoxifen), and EGFR inhibitors (cetuximab) ^4 5 6 7 8 21^. In our previous study ^15^ of investigating the relationship between p53 inactivation and chemotherapeutic resistance, we used NCI approved oncology drug set VI, consisting of 127 FDA approved anticancer drugs, to screen against TP53-WT and TP53-KO human embryonic stem cells (hESCs). We discovered that TP53-KO hESCs exhibited resistance to cisplatin, olaparib, and carboplatin, highlighting the challenge of treating cancers with these commonly used agents in the presence of TP53 mutations.

To establish our previous findings in human colon tumor organoids, we created and used an isogenic pair of TP53-WT and TP53-KO organoid (F147T) to investigate the relationship between p53 inactivation and resistance to cisplatin ^16^. We applied a simple high-throughput organoid plating method ^22^ of a 50/50 mixture of TP53-WT and TP53-KO organoids. Mixtures were then treated with seven different concentration of cisplatin and nutlin over a span of 10 days, refeeding every two days. After the treatment was completed, we prepared genomic DNA and used multi-color qPCR targeting unique sequences in the two different vectors to determine TP53-WT and TP53-KO quantities in each well. We found that both cisplatin and nutlin showed the expected selective effect on TP53-WT organoids. We determined the optimal concentration of cisplatin at which the TP53-WT colon tumor organoids were very sensitive, yet TP53-KO colon tumor organoids were very resistant (10 uM). These findings suggest that these colon cancer organoids depend on other genes for survival in the absence of functional TP53. Employing a genome-wide CRISPR KO library screen in the presence of cisplatin could help to identify these secondary genes, further elucidating the complex interactions that contribute to chemotherapy resistance.

### Genome-wide CRISPR/Cas9 screen with cisplatin for three-way synthetic lethal interaction

Three-way synthetic lethal interactions involving TP53 mutations, DNA-damaging chemotherapeutic agents, and a secondary gene mutation, are demonstrable. These secondary genes can be identified using genome-wide CRISPR knockout library screening conducted on TP53-KO derivative of human colon cancer organoids, both in the presence and absence of chemotherapy treatments.

We screened our TP53-KO colon tumor organoids, which were previously subjected to genome-wide CRISPR KO screening using quadruplicate transductions of the Brunello library ^16^ subsequently grown in the presence or absence of cisplatin to identify targets that re-sensitize these cisplatin-resistant colon organoids to this chemotherapy agent. The CRISPR library-screened organoids had been cultured for 58 days before starting the cisplatin treatment, which continued for 28 days (total of 86 days), with samplings at Day 58 (starting point), Day 65, Day 76, and Day 86 for both cisplatin and DMF (Fig. 1). At each time point, we prepared genomic DNA, PCR amplified the guides and sequenced the amplicons (BROAD Genetic Perturbation Platform ^23^). The abundance of each guide was determined by quantifying each unique gRNA sequence.

**Fig. 1:**
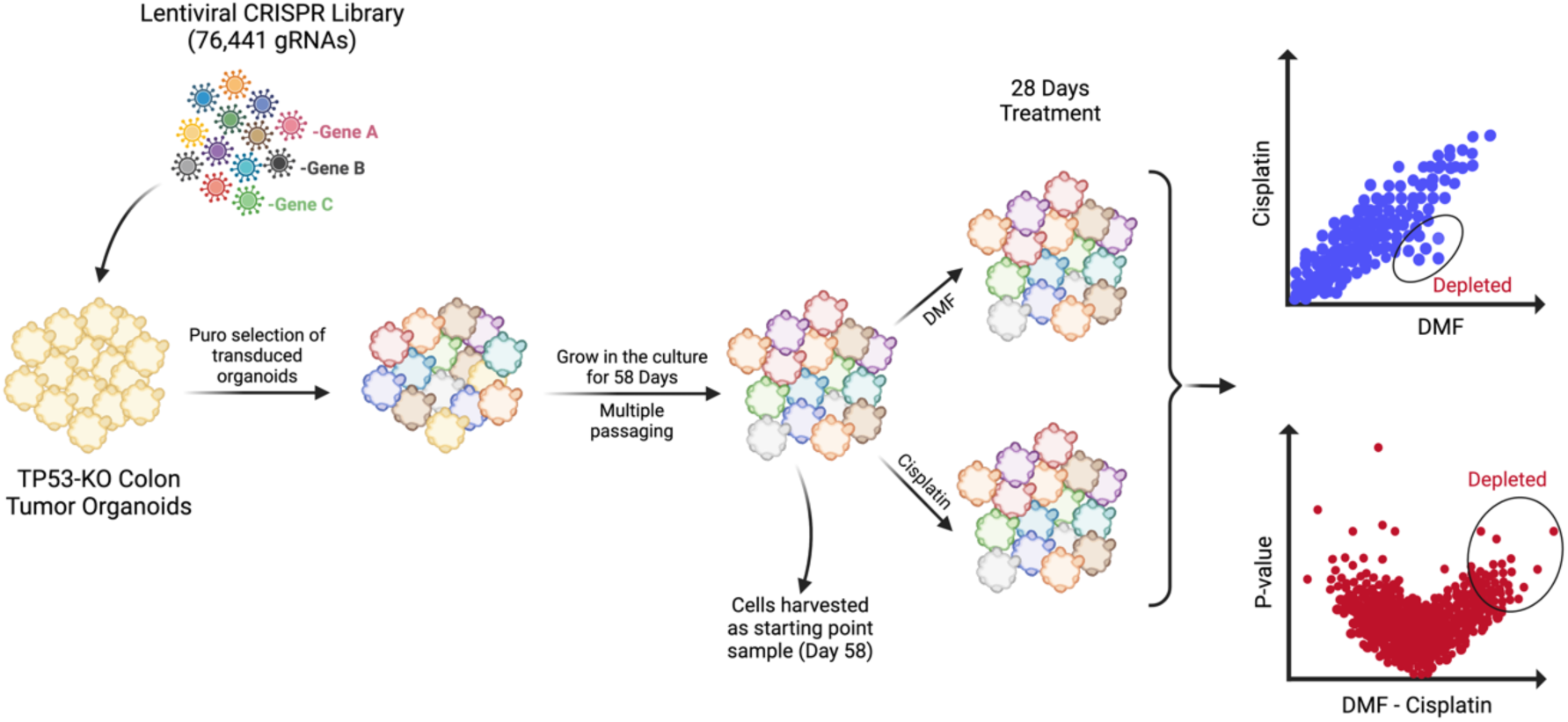
Genome-wide CRISPR knockout screen with cisplatin in TP53-KO colon cancer organoids. Workflow of the CRISPR screen in presence and absence of cisplatin. Created with BioRender.com

Fig. 2A shows the distribution of log-normalized gRNA frequencies for cisplatin treated and no drug (DMF) control samples and Fig. 2B shows the significance of the differences between the two classes. Guides with substantial shift to the right quadrant of the volcano plot had a greater mean difference and higher significance (p-value <0.05) in their impact on cell viability post-cisplatin treatment (Fig. 2B). We observed most of the guides significantly depleted in cisplatin samples compared to DMF, correspond to a set of genes involved in the DNA repair pathways including Fanconi Anemia (FA) (p-value <0.05). Enrichment of these genes was identified through Gene Set Enrichment Analysis (GSEA) ^24^ (Fig. 3A and Table1). Targeting these pathways could enhance the sensitivity of TP53-mutant colon cancer cells to cisplatin (Fig. 3B). The genes in these pathways are also expressed in normal colon tissues, confirmed through the BROAD Adult Genotype Expression (GTEx) database (data not shown). Among those genes enriched, FANCL, BRIP1 (FANJ), and ERCC6 are top ranked (p-value <0.05), and they could be potential synthetic lethal targets for cisplatin-resistance TP53 mutant colon tumors.

**Fig. 2:**
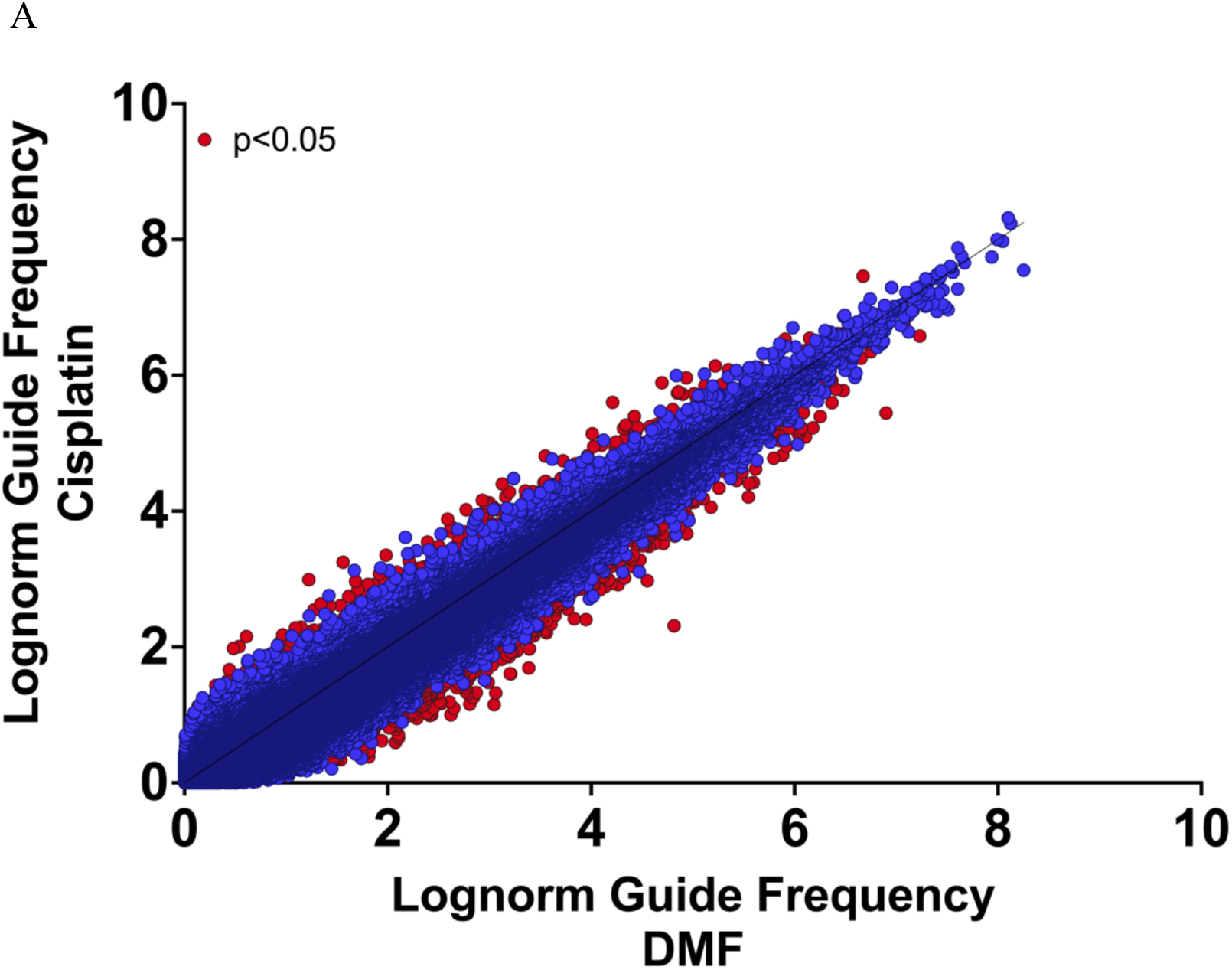
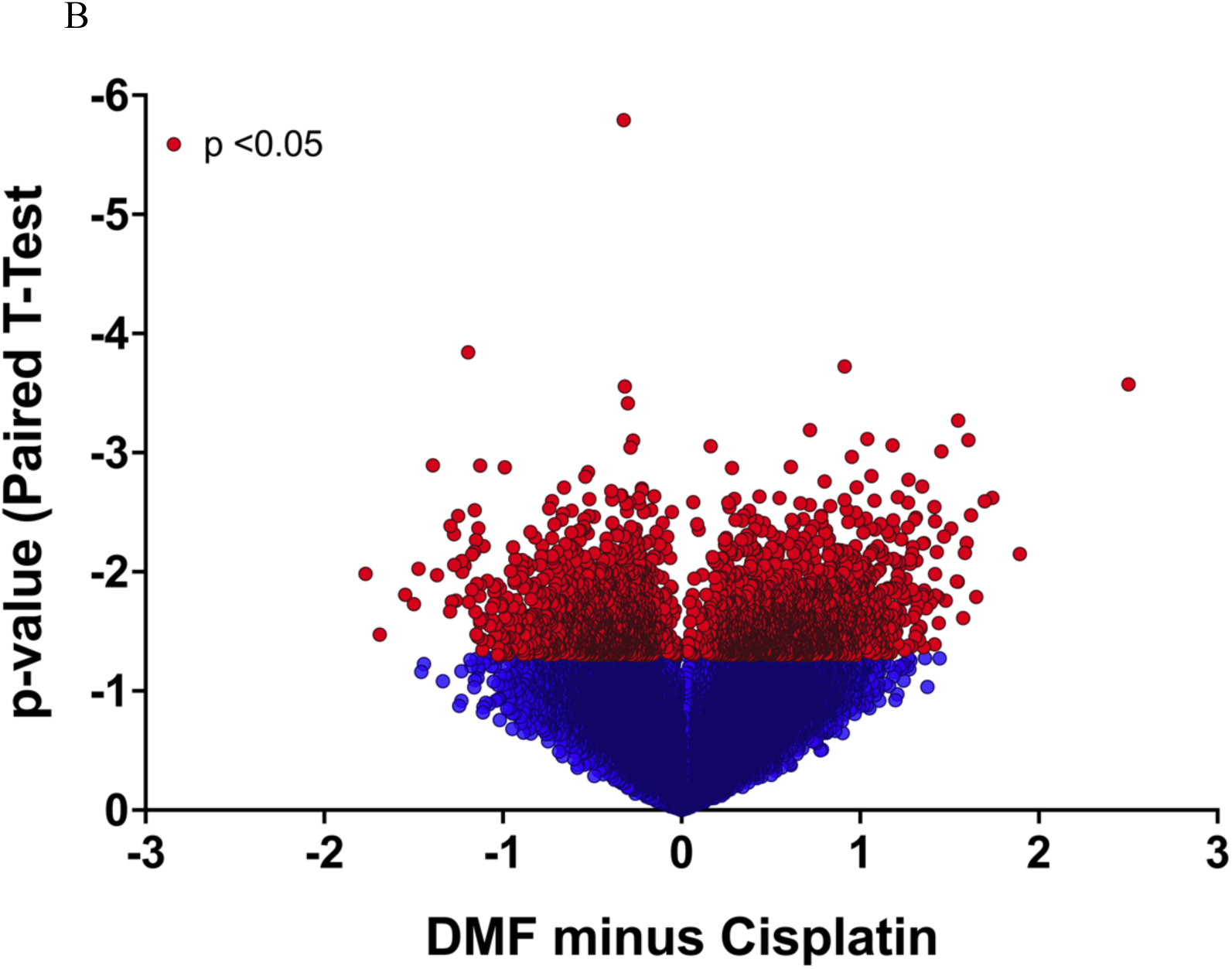
gRNA distributions. (**A**) Distribution of average log-normalized gRNA frequencies in TP53-KO organoids treated with cisplatin compared to no drug (DMF). The red points indicate guides significantly different between the two conditions (paired t-test unadjusted P<0.05). (**B**) Volcano plot illustrating the statistical significance and mean difference in gRNA effects on cell viability between cisplatin treatment and no drug (DMF). Significant guides on the right were depleted in presence of cisplatin.

**Fig. 3:**
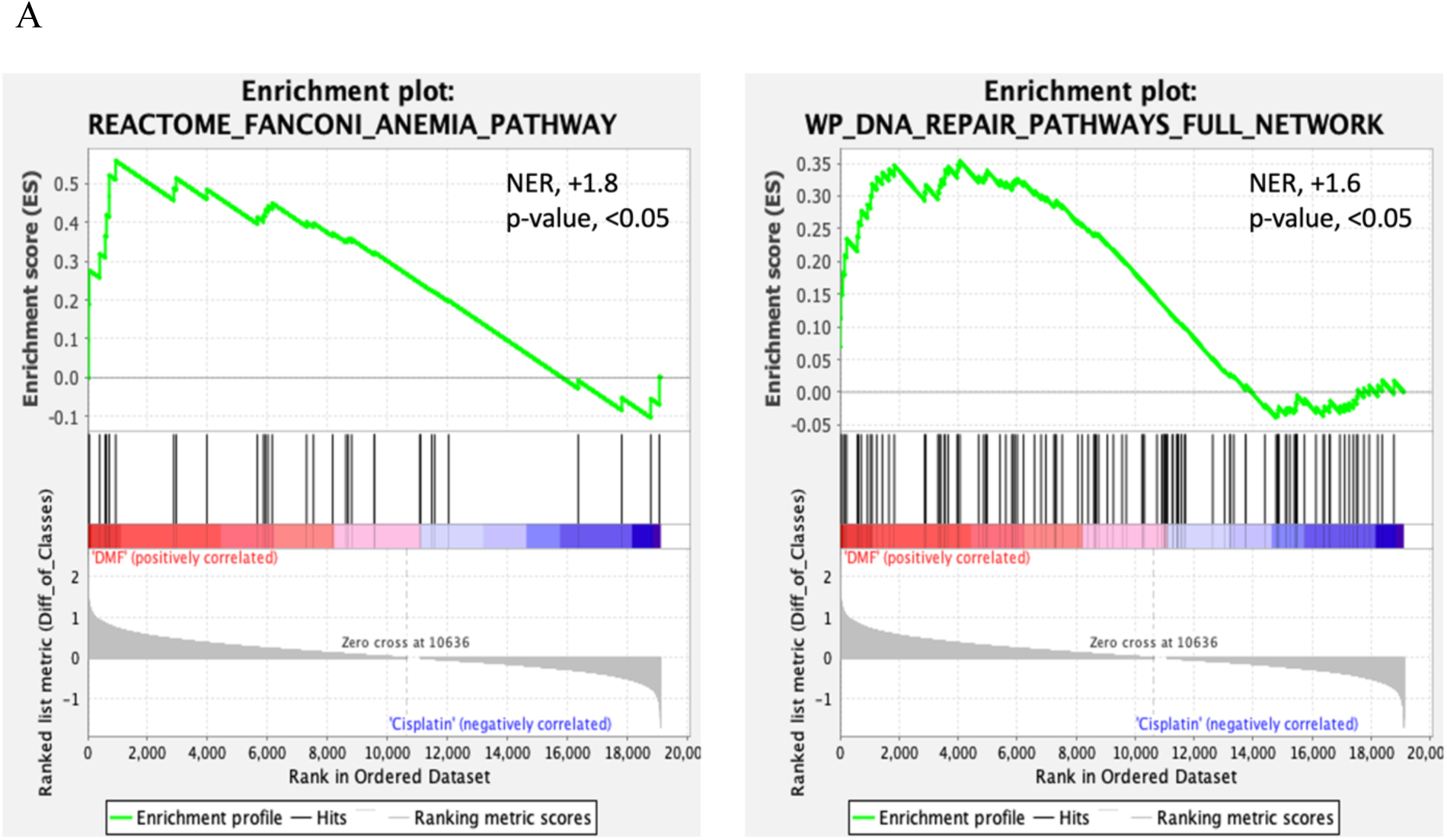
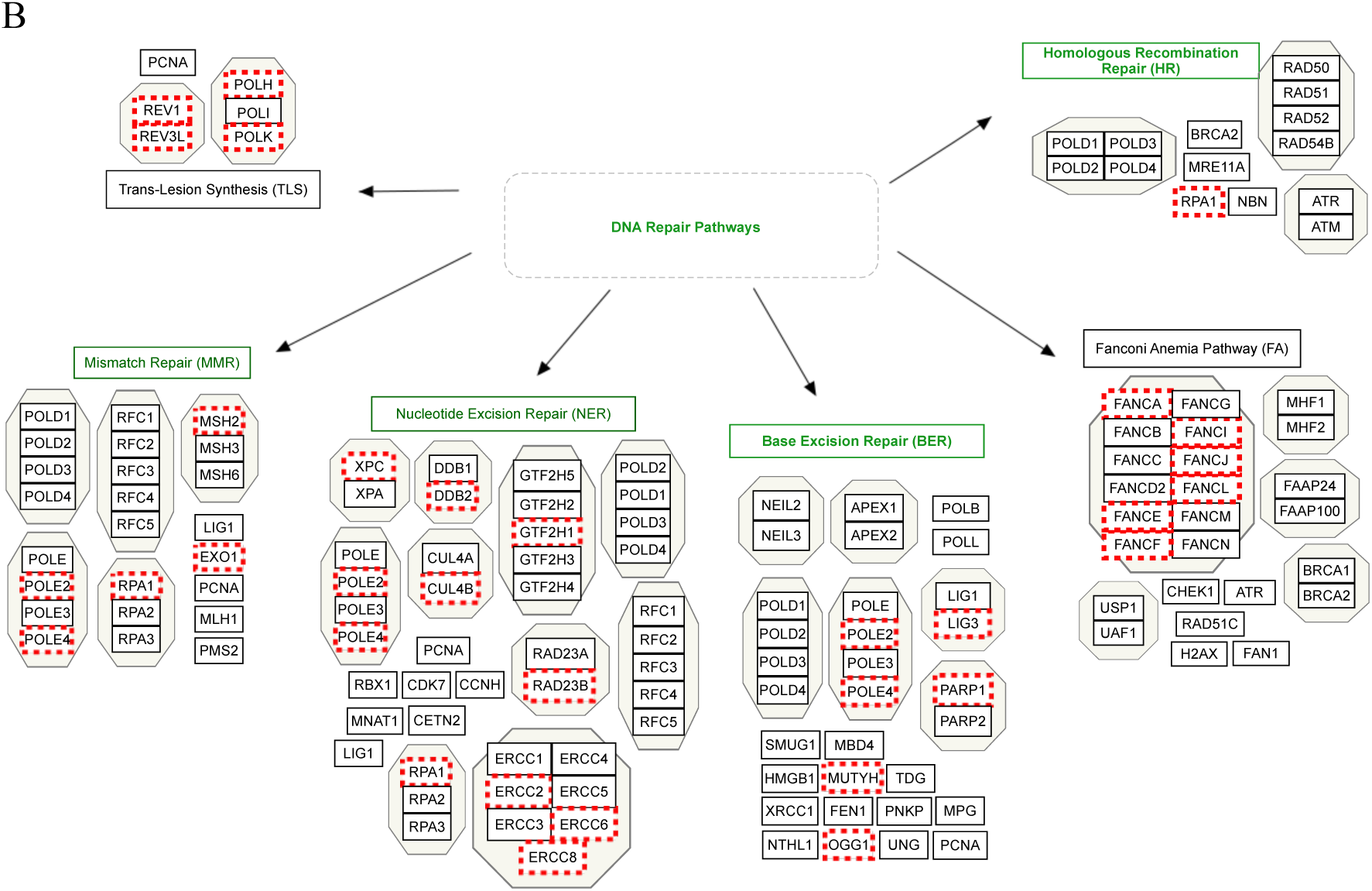
Gene Set Enrichment Analysis (GSEA). **(A)** Gene Set Enrichment Analysis reveals enrichment of depleted gRNAs targeting genes in the Fanconi Anemia (FA) and other DNA repair pathways. **(B)** The full network diagram of DNA Repair Pathways. Genes labeled with red are depleted in presence of cisplatin and are potential candidates for three-way synthetic lethal interaction. The diagram is adapted from DAVID ^35 36^.

**Table. 1.**
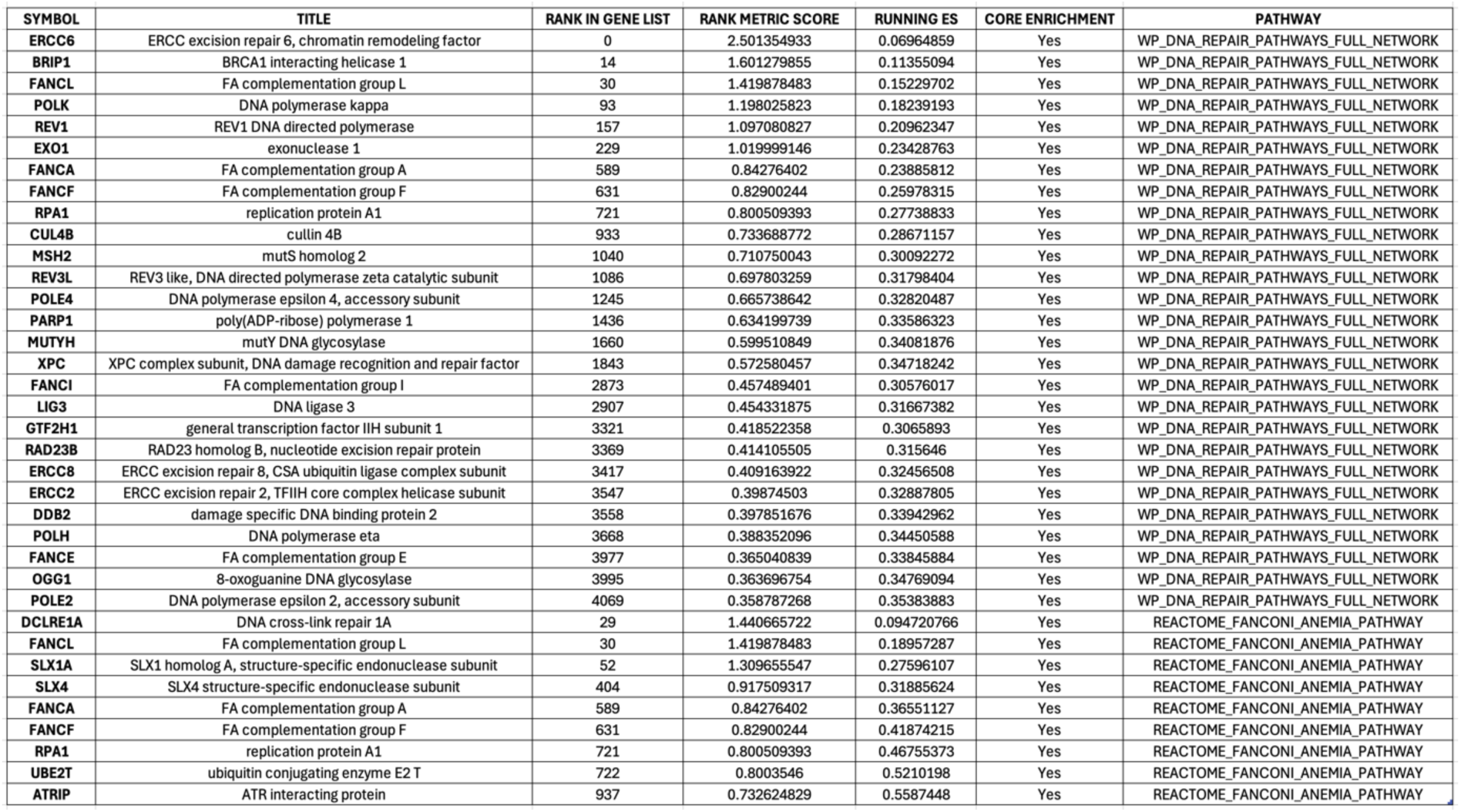
The Enriched gene list in DNA repair (top) and FA (bottom) pathways. FANCL, ERCC6, and BRIP1 are top-ranked depleted guides targeting genes in these pathways.

Fig. 4 shows how the log-fold changes (LFCs) of the gRNAs (relative to start of cisplatin treatment) changes over time with cisplatin treatment. Guides corresponding to the Faconi Anemai (FA) and DNA repair genes identified as significantly enriched by GSEA are highlighted and are more depleted at later time points in the cisplatin treatment compared to DMF control. The changes in the abundance of the significant gRNAs over time from select genes of the DNA repair pathways are shown in dot plots (Fig. 5). As expected, TP53-KO organoids are resistant to MDM2 knockout in presence and absence of cisplatin. However, the depletion of ERCC6, FANCL, and BRIP1 gRNAs result in a more pronounced reduction in organoid viability in cisplatin treated compared to the DMF controls.

**Fig. 4:**
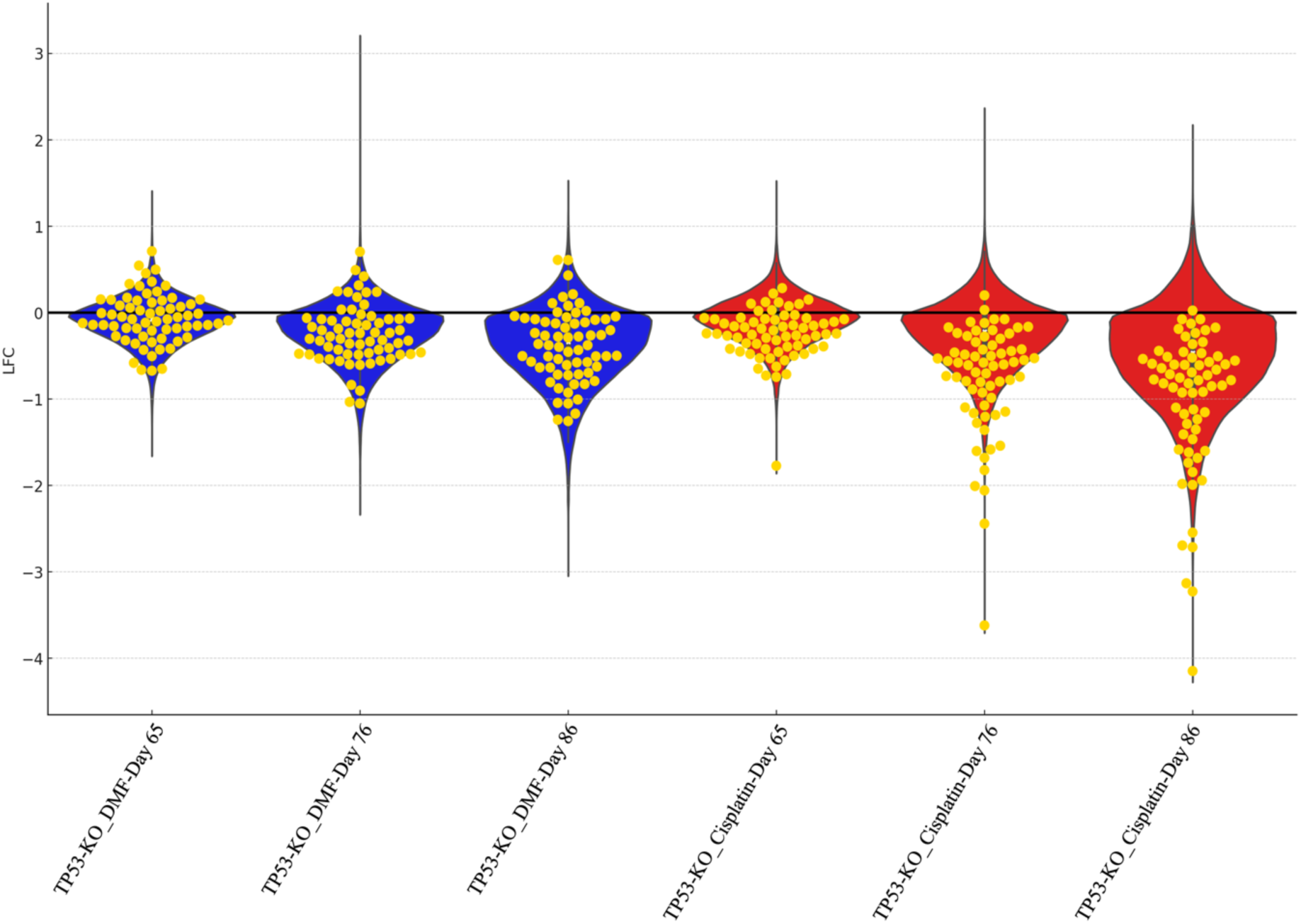
Log fold changes of gRNAs in TP53-KO colon cancer organoids treated with cisplatin and DMF. Distribution of average gRNA log-fold changes over time comparing DMF and cisplatin treatments to day 58 (before starting the cisplatin treatment). Golden dots are gRNAs targeting genes in Table 1. Depletion under cisplatin treatment suggesting these genes are needed for resistance to cisplatin.

**Fig. 5:**
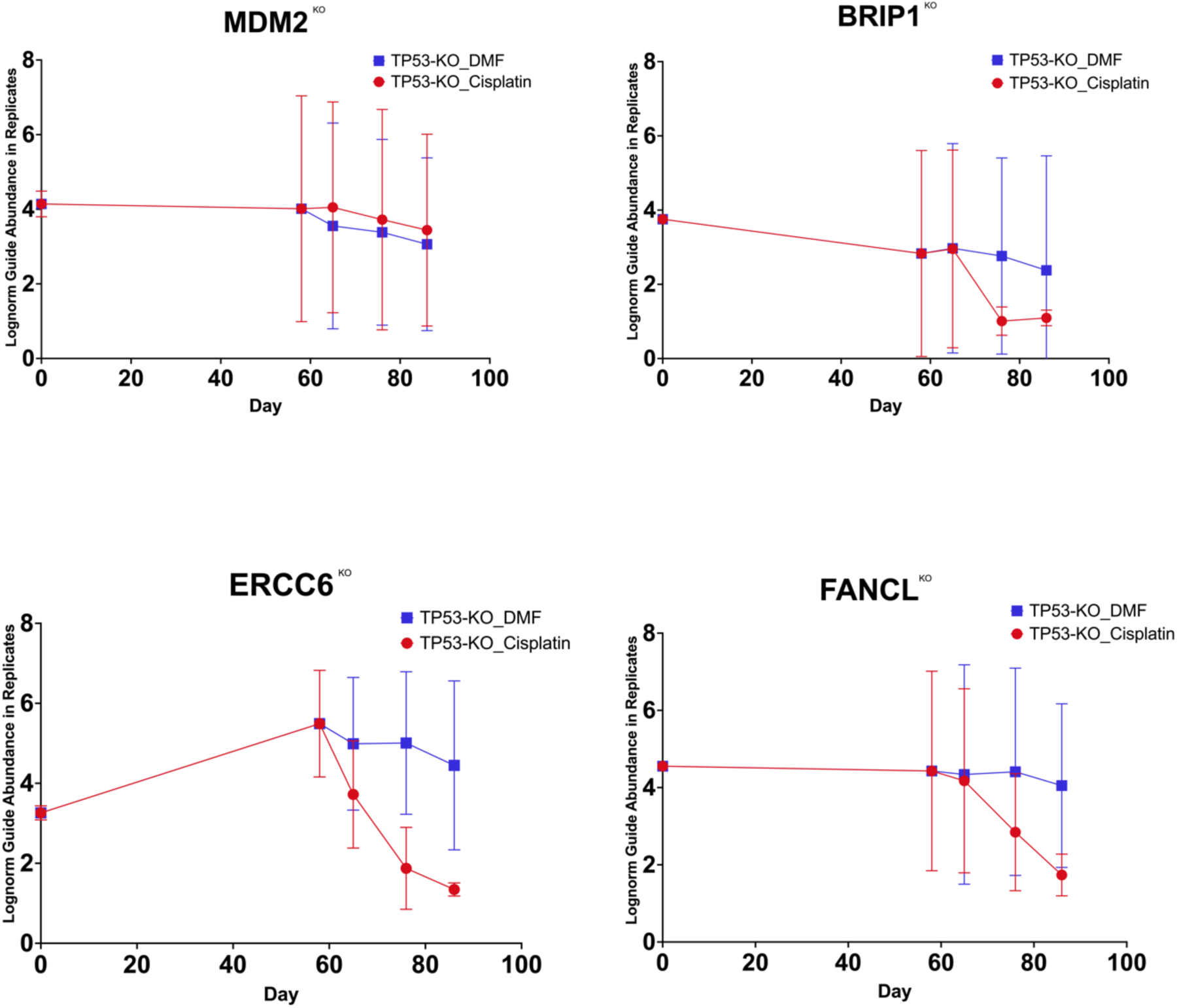
CRISPR dropout screen with cisplatin in TP53-KO colon cancer organoids. Dot plot of select significantly depleted guides (p-value<0.05). As expected, TP53-KO organoids are resistant to MDM2 knockout in presence and absence of cisplatin. However, the depletion of ERCC6, FANCL, and BRIP1 gRNAs result in a more pronounced reduction in organoid viability in cisplatin treated compared to the DMF controls.

It can be hypothesized that, in the absence of functional TP53, tumor cells exposed to DNA damaging agents become increasingly reliant on alternative repair pathways, such as the Fanconi Anemia (FA), nucleotide excision repair (NER), and mismatch repair (MMR) pathways, to repair DNA damage and maintain survival. The activities of these pathways enables tumor cells to withstand genotoxic stress induced by DNA-damaging agents like cisplatin, thereby contributing to cisplatin resistance ^19 25 26^. Among these pathways, the FA pathways play a central role in repairing interstrand cross-links (ICLs), one of the primary forms of DNA damage induced by cisplatin. The FA core complex, an E3 ligase composed of nine distinct proteins ^27 28^, is essential for monoubiquitination of FANCD2/FANCI. This modification activates downstream repair proteins, facilitating the resolution of intrastrand cross-links ^27 28 25^. Similarly, ERCC6, a key protein in the transcription-coupled repair sub-pathway of nucleotide excision repair, plays a crucial role in resolving DNA lesions on actively transcribed genes ^29 30^. Its efficient function in repairing cisplatin-induced cross-links contributes to chemoresistance by allowing tumor cells to survive despite ongoing DNA damage. These dual roles of DNA repair proteins like ERCC6 highlight their potential as therapeutic targets in TP53-mutant tumors.

### Validation

To validate select candidate genes, we tested select candidates, ERCC6, FANCL, and BRIP1, we first created stable KOs of each gene in TP53-KO organoid (F147T) using lentivirus carrying single gRNAs targeting each gene, along with a non-targeting control guide, selected with puromycin for 7 days. We then created a mixture of these stable KOs in triplicates and cultured in presence and absence of cisplatin for 27 days. Time point samples were taken on Day 0 (starting point of cisplatin treatment), Day 9, Day 15, Day 21 and Day 27. At each time point, we collected cells and prepared genomic DNA and counted the guides using PCR and Nanopore amplicon sequencing. The gRNA depletion was determined by counting unique gRNA sequences and normalizing these counts to the counts of non-targeting control guide within each sample (frequency). Results, as shown in Figure 6, validated three-way synthetic lethal interactions of ERCC6, FANCL, or BRIP1 with TP53-KO and cisplatin treatment. Knockout of either of these genes effectively results in re-sensitization of the TP53-KO organoids to cisplatin. These findings demonstrate that ERCC6, FANCL, and BRIP1 depletion enhances the anti-tumor effect of cisplatin *in vitro*. The entire group of enriched genes identified through GSEA are promising candidates for drug development as potential strategies to overcome drug resistance or improve drug efficacy.

**Fig. 6:**
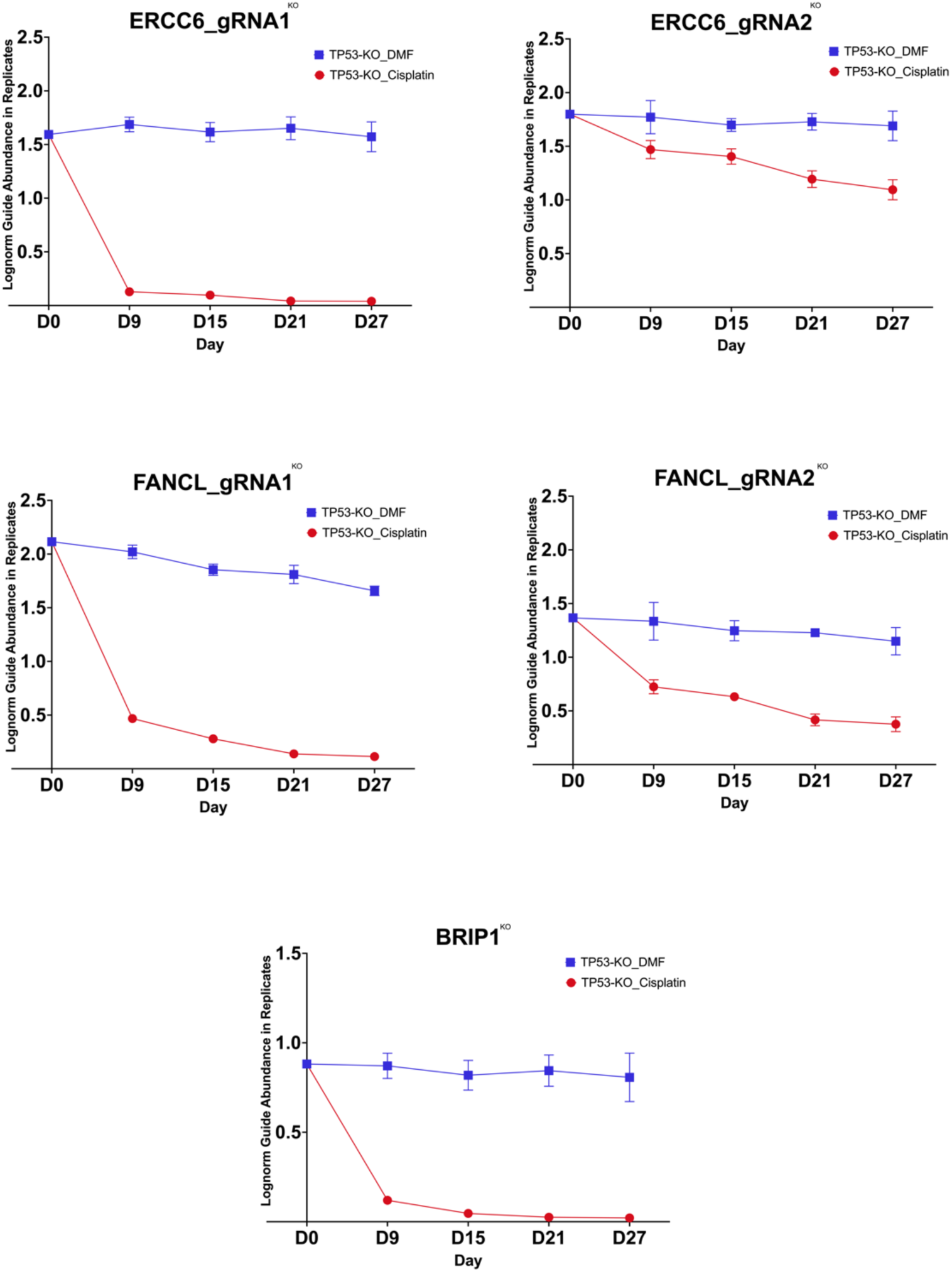
Validation of select targets. Dot plot of validated guides show that depletion of ERCC6, FANCL, or BRIP1 gRNAs result in pronounced sensitization of TP53-KO organoids to cisplatin.

## Discussion

In this work, we investigated the underlying mechanisms of chemoresistance in TP53-null colon cancer organoids and identified potential targets to resensitize these cells to cisplatin therapy. TP53, often known as the “guardian of the genome,” plays a pivotal role in DNA damage response ^4^, and its inactivation, common in colorectal cancer, is linked to impaired apoptosis and resistance to DNA-damaging agents like cisplatin ^19 20 21^. The observed resistance in TP53-KO colon tumor organoids emphasizes the critical role of tumor suppressor pathways in chemosensitivity and aligns with prior studies highlighting the link between TP53 mutations and chemoresistance ^4 5 6 7 8 19 20 21^. The identification of secondary genes through CRISPR/Cas9 screenings offers a promising avenue for the development of targeted therapies that could be used in combination with existing chemotherapeutics ^10^. The three-way synthetic lethal interactions uncovered in this study suggest that targeting secondary pathways could lead to a more robust apoptotic response, overcoming the cisplatin resistance often seen in TP53-mutant tumors. This approach could refine the use of cisplatin, potentially reducing the required dosage and associated toxicities.

Our CRISPR/Cas9 library screen on human colon cancer organoids identified essential genes within the DNA Repair pathways, including Fanconi Anemia (FA) pathway. Among these, FACNL, BRIP1 (FANLJ), and ERCC6 emerged as potential therapeutic targets to enhance cisplatin efficacy. These pathways play a crucial role in repairing cisplatin-induced DNA cross-links ^19 25 26 31 32 33 27 34^, and their inhibition introduces synthetic lethality in TP53-null cells. This study advances our understanding of cisplatin resistance in TP53-null colon cancer. Targeting the DNA repair and FA pathways could re-sensitize these cancers to treatment and improve therapeutic outcomes. Our findings highlight the potential of leveraging genetic screening to identify vulnerabilities in cancer cells, paving the way for more effective therapeutic strategies. Further exploration through *in vivo* studies and clinical trials can strengthen the translational impact of these findings, with the potential to extend their applicability to other TP53-mutant cancers.

## Data Availability

All datasets generated and analyzed during the study are available from the corresponding authors on reasonable request.

## Methods

### Organoid Isolation and Culture

Organoids were generated from de-identified / anonymized surgically resected colorectal cancers ^37^. Small pieces (1cm square) of mucosal tissue were separated and washed in 1X PBS (Corning) supplemented with 0.1% Amphotericin B (Gibco) and 0.2% Primocin (Invivogen) to remove bacterial. Crypts were isolated with digestion and incubation in 1X PBS supplemented with 10% Collagenase A (Sigma) and 10 μM Y-27632 RhoKinase inhibitor (Selleckchem) on shaker at 37°C for 40 min. following with adding 5% FBS (Gibco), and incubation for fewer min at room temp. Crypts were dislodged mechanically from tissue by pipetting up and down, separated into a new tube for additional wash and were centrifuged at 500 x g for 5 min. The isolated stem cells were embedded in 75ul of ice-cold RGF BME Matrigel (R&D systems) per well in 12-well plates. The Matrigel was polymerized for 10-15 minutes at 37°C humidity-controlled incubator with 5% CO2 and covered with 2ml per well of human colon organoids media (ENRA) containing 10 μM Y-27632 RhoKinase inhibitor (Selleckchem. The ENRA media contained Advanced DMEM/F-12 media (Gibco) supplemented with 10% R-Spondin, 10% Noggin (all CM produced in-house), 1x N2, 1x B27, 10 mM HEPES, 1x Glutamax, 1% Penicillin/Streptomycin, 0.25 ugmL^−1^ Amphotericin B, 50 ngmL^−1^ human Epidermal Growth Factor EGF (all from Gibco), 10 mM Nicotinamide, 1.25 mM N-acetyl-cysteine, 10 nM Gastrin (all from Sigma), 0.5 μM TGF-β type I receptor inhibitor A83-01 (Tocris Bioscience), 100 ugmL^−1^ Primocin (Invivogen). Media was changed every other day. For the first two changes of media after culturing, media contained 10 μM Y-27632 RhoKinase inhibitor (Selleckchem). Organoids were passaged 1:4 every 10-14 days. For passaging, organoids and matrigel were covered with 2ml/well of TrypLE (Gibco) plus 10 μM ROCK inhibitor (Selleckchem), dissociated by incubation at 37°C for 5-7 min, then spun down at 500g for 5 min and were mechanically disrupted using a P1000 pipette and transferred into a 15-ml conical tube. Centrifuged at 500 x g for 5 min. The pellet was resuspended and plated in ice-cold fresh Matrigel (R&D systems) and covered with ENRA media. Organoid lines were constantly tested for mycoplasma contamination and resulted negative.

### Cell Culture

All colon tumor organoids were grown as described before ^37^. Organoids were maintained in ENRA media containing 1% Penicillin/Streptomycin, 100 μgmL^−1^ Primocin, 0.25 ugmL^−1^ Amphotericin B. Organoids were kept in a 37°C humidity-controlled incubator with 5% CO2 and were maintained in exponential phase growth by passaging every 10-14 days and media was changed every other day. For viral transduction, 6μgmL^−1^polybrene was added during spinfection. After viral transduction, 2μgmL^−1^ puromycin or 20μgmL^−1^ blasticidin were used for selection.

### Vectors

Individual gRNA sequences are provided in Supplementary Table 1 from the Brunello library and select candidate guide sequences are provided in Supplementary Table 1 used for validation. The following vectors were used in the study ^16^ and are available on Addgene:

pRDA_186 (Spyo-only anchor vector): U6 promoter expresses customizable Spyo-guide; PGK promoter expresses blasticidin resistance and 2A site provides EGFP (Addgene 133458).

lentiCRISPRv2 (pXPR_023): EF1a promoter expresses SpyoCas9 and 2A site provides puromycin resistance; U6 promoter expresses customizable Spyo-guide (Addgene 52961).

### Viral Transduction

Prior to viral transduction, organoids were digested to dissociate them into single cells by incubation in TrypLE contained 10 μM ROCK inhibitor at 37°C for 20-30 min. Then, the cell suspension was passed through a 40um Nylon Mesh cell strainer to segregate any cell clumps present. Following this, cells were counted to separate the required number of cells for transduction and resuspended in 1X human colon tumor organoids media (ENRA) contained 10μM ROCKi and Polybrene (6μg mL^−1^). Next, the relevant viral particles, determined based on the desired multiplicity of infection (MOI), were added to the cell mixture. The mixture was then transferred to a tissue culture plate, and it was spun at 1000 x g for 2 h at 30°C. Post-spinfection, the plate incubated at 37°C incubator with 5% CO2 for additional 3 h (total of 5 h). Following the incubation period, virus was washed out by ice-cold PBS contained 10μM ROCKi from the organoids. Organoids then were embedded in ice-cold Matrigel and covered with ENRA media contained 10 μM ROCKi inhibitor for the first two media changes. For all the other feedings, the organoids were maintained in ENRA media without ROCKi. The selection was started 2 days after the viral transduction.

### Brunello CRIPSR Library Screening in Presence and Absence Cisplatin

Brunello library was purchased from Addgene (73179-LV). Generating TP53 anchor vector derivative and transduction for Brunello library screen in quadruplicates was performed as before ^16^. Throughout the screen, cells were plated into Matrigel domes in 150mm dishes. Replicates and each treatment condition were kept separate through the whole screen. At each sampling time point, organoids were split at a density to maintain a representation of at least 200X for each replicate and cell counts were taken at each sampling time point to monitor growth. A sample was taken for all replicates at Day 58 prior to starting the cisplatin treatment as the baseline, then Day 65, Day 76, and Day 86. For each sampling, organoids were digested to single cell in TrypLE at 37°C bead bath for 30 min, pelleted by centrifugation, resuspended in PBS and divided in half. One half was plated back in Matrigel to grow for later time points and the other half was used for genomic DNA isolation.

### Validation

lentiCRISPR single gRNA clones and lentiviruses were produced by VectorBuilder. Throughout the validation mixture experiment, cells were plated into Matrigel domes in 6-well plates. Replicates and each treatment condition were kept separate through the whole experiment. At each sampling time point, organoids were split at a density to maintain a representation of at least 1000X for each replicate and cell counts were taken at each sampling time point to monitor growth. A sample was taken for all replicates at Day 0, when the mixture was created, prior to starting the cisplatin treatment as the baseline, then Day 9, Day 15, Day 21, and Day 27. For each sampling, organoids were digested to single cell in TrypLE at 37°C bead bath for 20-30 min, pelleted by centrifugation, resuspended in PBS and divided in half. One half was plated back in Matrigel to grow for later time points and the other half was used for genomic DNA isolation.

### Genomic DNA isolation

Genomic DNA (gDNA) was isolated using the Blood & Cell Culture DNA Midi Kits (Qiagen) as per the manufacturer’s instructions. The gDNA concentrations were quantitated by Qubit.

### Sequencing

The gDNA of Brunello samples, treated with cisplatin and DMF, were PCR amplified with primers that flank the Gecko guide RNA inserts sequences (Gecko_F and Gecko_R) and sequenced the PCR products by illumine platform (BROAD Genomics Perturbation Platform). The representation of at least 200X coverage was kept. For validations, gDNA was PCR amplified with primers that flank the Gecko guide RNA inserts sequences (Gecko_F and Gecko_R) and sequenced by Oxford Nanopore platform in-house using the NBD112.24 (Q20+) kit and kept a representation of at least 1000X coverage. All reads were counted by aligning the sequence reads to a reference file of all possible gRNAs present in libraries. This process made it possible to count all the gRNA inserts, setting the stage for further detailed examination and analysis.

Gecko primers used for Brunello library amplification:

Forward primer, 5′–3′; TCTTGTGGAAAGGACGAAACACCG

Reverse primer, 5′–3′; TGTGGGCGATGTGCGCTCTG

### Sample Sizes and Statistical Techniques

Brunello CRISPR dropout screen with cisplatin and DMF was performed in quadruplicates for all time points.

Validation mixture experiment with cisplatin and DMF was performed in triplicates for all time points.

Dots plots display the mean ± SD from replicates (four replicates in the Brunello screen and three replicates in the validation).

### Statistical Analysis of the Brunello Screen with Cisplatin

Gene set enrichment analysis was performed with the local version of the GSEA tool http://www.broadinstitute.org/gsea/index.jsp. Log-normalized counts of unique gRNA sequence from ∼78,000 guides in the Brunello library were used with collapsing to gene symbol and max probe mode. All gene sets, and genes were ranked according to different of classes.

To determine the abundance of each guide, we calculated the average of log-normalized counts for sample treated with cisplatin and those with DMF to facilitate a comparative analysis between the two-treatment group (Fig 2a and supplementary Table.2).

To determine the statistical significance of each gRNA, we used the average of log-normalized frequencies to calculate the mean difference between DMF and cisplatin samples for each gRNA. To assess the significance of the difference between the two conditions, we performed a T-test comparing the normalized frequencies in sample treated with cisplatin and those treated with DMF. Guides with *P*-value smaller than 0.05 are highlighted in red in Fig. 2b (supplementary Table.2)

To visualize the distribution of gRNAs over time, we calculated the log fold change (LFC) for each time point by subtracting the log-normalized counts of each unique guide in each replicate from its initial time point counts (Day 58, the point when the TP53-KO pool of library-infected organoids was divided for the DMF and cisplatin treatments) (Fig.4).

### Statistical Analysis of the Validation

Following deconvolution, the abundance of each guide was determined by unique gRNA sequence counting (oxford nanopore ligation sequencing of gRNA amplicon – NBD112.24 (Q20+)) and normalizing these counts to the average counts from all guide sequences within each sample. We conducted a comparative study of the gRNA depletion between cisplatin samples and DMF samples.

### Data Visualization

Scientific illustration was generated with Biorender.com. Graphs were generated using GraphPad Prism.

## Acknowledgement

This study was supported by the US Department of Defense CDMRP PRCRP Grant CA180925. Organoid biobanking was supported by an NCI SBIR to Dr. Carolyn Banister.

## Ethics Statement

All samples were collected under an IRB approval protocol (Pro00022064) “The Palmetto Health – University of South Carolina Biorepository”.

## Author Contributions

S.K, and P.J.B designed all the experiments. S.K and P.H carried out all experimental work. S.K and P.J.B designed and carried out all computational analysis. P.H and M.K contributed to the performance of drug screen experiments. C.E.B established colon human tumor’s organoid biobank. S.E.M provided colon human tumor tissues. P.J.B supervised the work. S.K and P.J.B wrote the manuscript, with input from all the authors.

Supplementary Information is available for this paper.

## Supplementary Information

**Supplementary Table.1**

Quantitative analysis of gRNA depletion in CRISPR-Cas9 library screen in presence and absence of cisplatin. Guide RNA (gRNA) abundance was assessed by counting unique gRNA sequences and normalizing the counts against the average counts from a control set of 1,000 non-targeting gRNAs. The average of the log-normalized count for sample treated with cisplatin and those with DMF was calculated to facilitate comparative analysis between the two-treatment groups. A T-test comparing the normalized frequencies in sample treated with cisplatin and those with DMF was calculated. The mean difference between DMF and cisplatin samples for each gRNAs was calculated using the average of log-normalized frequencies.

**Supplementary Table.2**

gRNA sequences used for validation study.

